# TAPIR: a T-cell receptor language model for predicting rare and novel targets

**DOI:** 10.1101/2023.09.12.557285

**Authors:** Ethan Fast, Manjima Dhar, Binbin Chen

**Author notes:** **Corresponding authors:** Ethan Fast, Binbin Chen.

## Abstract

T-cell receptors (TCRs) are involved in most human diseases, but linking their sequences with their targets remains an unsolved grand challenge in the field. In this study, we present TAPIR (T-cell receptor and Peptide Interaction Recognizer), a T-cell receptor (TCR) language model that predicts TCR-target interactions, with a focus on novel and rare targets. TAPIR employs deep convolutional neural network (CNN) encoders to process TCR and target sequences across flexible representations (e.g., beta-chain only, unknown MHC allele, etc.) and learns patterns of interactivity via several training tasks. This flexibility allows TAPIR to train on more than 50k either paired (alpha and beta chain) or unpaired TCRs (just alpha or beta chain) from public and proprietary databases against 1933 unique targets. TAPIR demonstrates state-of-the-art performance when predicting TCR interactivity against common benchmark targets and is the first method to demonstrate strong performance when predicting TCR interactivity against novel targets, where no examples are provided in training. TAPIR is also capable of predicting TCR interaction against MHC alleles in the absence of target information. Leveraging these capabilities, we apply TAPIR to cancer patient TCR repertoires and identify and validate a novel and potent anti-cancer T-cell receptor against a shared cancer neoantigen target (PIK3CA H1047L). We further show how TAPIR, when extended with a generative neural network, is capable of directly designing T-cell receptor sequences that interact with a target of interest.

## Introduction

The adaptive immune system mobilizes a vast array of T-cells with diverse T-cell receptors (TCRs) to recognize and respond to diverse targets in cancer, infections, and autoimmune disease^1–5^. T-cell receptors are estimated to recognize up to billions of different targets^6^ and have been described as a fundamental language of the human immune system^7^. However, the interaction between TCRs and their targets remains largely enigmatic, characterized by a complex interplay between a TCR’s alpha and beta chains, an MHC protein, and a target peptide^1,8,9^.

In recent years, the development of machine learning based tools for antigen and TCR prediction has emerged as a promising avenue for improving TCR-based diagnostics^10,11^, cell therapies^12^, and vaccine design^13–17^. While existing tools have demonstrated strong performance on well-characterized targets (e.g., influenza or CMV), the field has struggled to make useful predictions about rare or novel targets such as mutated viral protein and cancer-associated neoantigens^18–20^. In this paper, we present a TCR language model that can predict TCR-target interaction for rare and novel targets never encountered in training.

Training machine learning models to generalize across millions of potential TCRs and thousands of potential targets requires advances in both model architecture and data generation. Inspired by the versatility of modern large language models, we developed TAPIR (<u>T</u>-cell receptor <u>a</u>nd <u>P</u>eptide Interaction <u>R</u>ecognizer), a deep neural network architecture that addresses these challenges. TAPIR leverages convolutional neural network based encoders to process TCR and target sequences across a variety of representations and several training tasks, predicting interactivity between sequences (**Fig. 1**). Resulting TAPIR models can make predictions over any combination of V gene, J gene, CDR3 gene, and MHC allele and target sequences, including sequences with missing information. This flexibility is based on TAPIR’s ability to learn from larger and more diverse datasets than prior work (**Fig. 2a**, **Fig 3a**), including a dataset of 25,207 paired and 21,180 unpaired TCRs against 1063 targets from public databases^5,21,22^, as well as a proprietary dataset of 4059 paired TCRs against an additional 932 targets (**Fig. 4a**).

**Figure 1.**
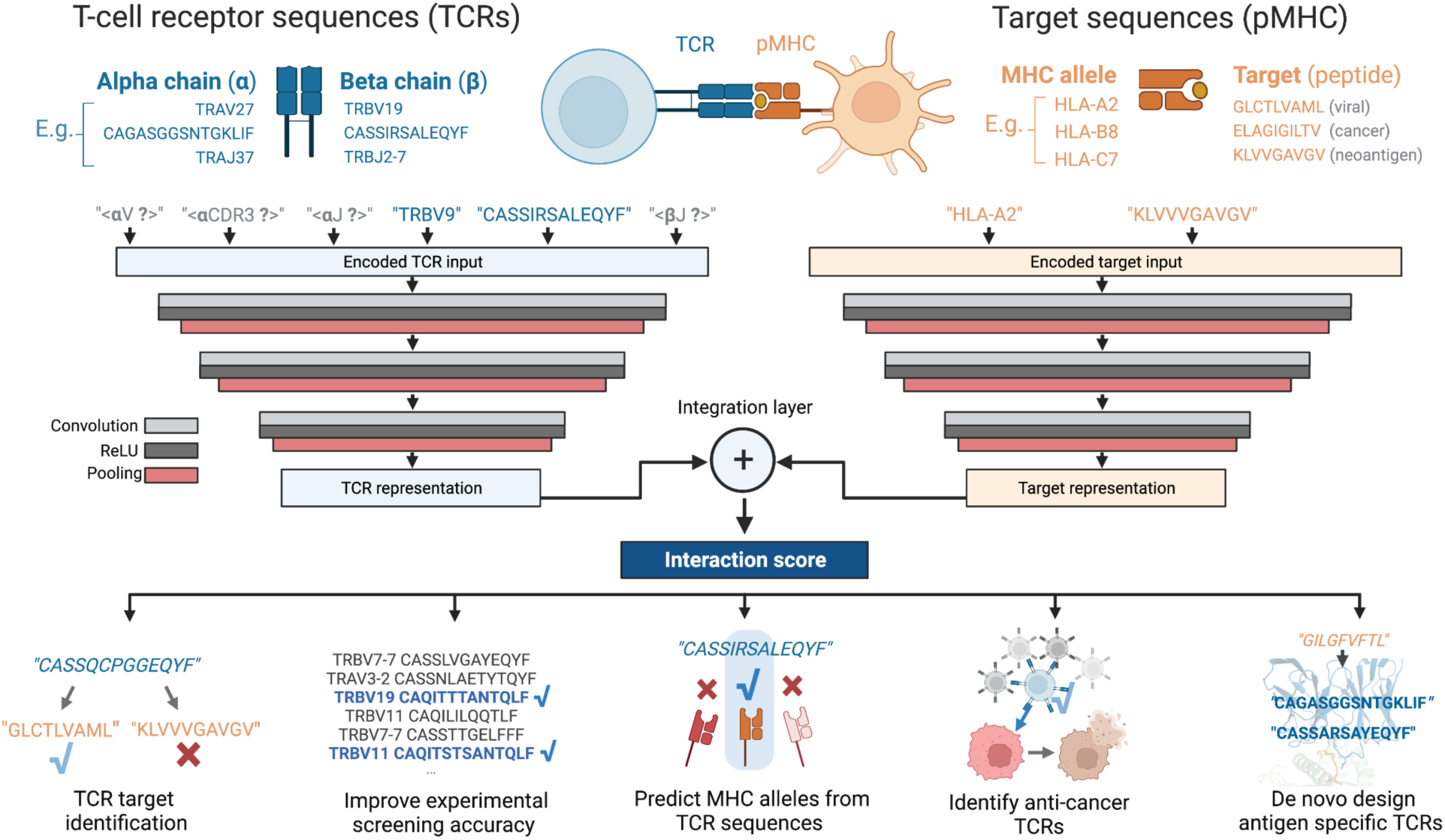
TAPIR: a T-cell receptor language model for predicting rare and novel targets. TAPIR takes input TCR and target sequences across any representation (e.g., beta chain only, paired chain but CDR3 only, etc.) and outputs an interaction score. TCRs and targets are encoded as sequences of amino acids and passed through a series of convolutional layers to create learned feature representations, which are then evaluated for interactivity through a final dense layer for classification. TAPIR’s architecture allows the model to predict TCR interactivity against novel targets that never appeared in its training data, such as cancer neoantigens with no known interacting TCRs. We demonstrate applications of TAPIR in improving experimental screening, in-silico TCR design, and identifying a novel TCR against a shared cancer neoantigen.

**Figure 2:**
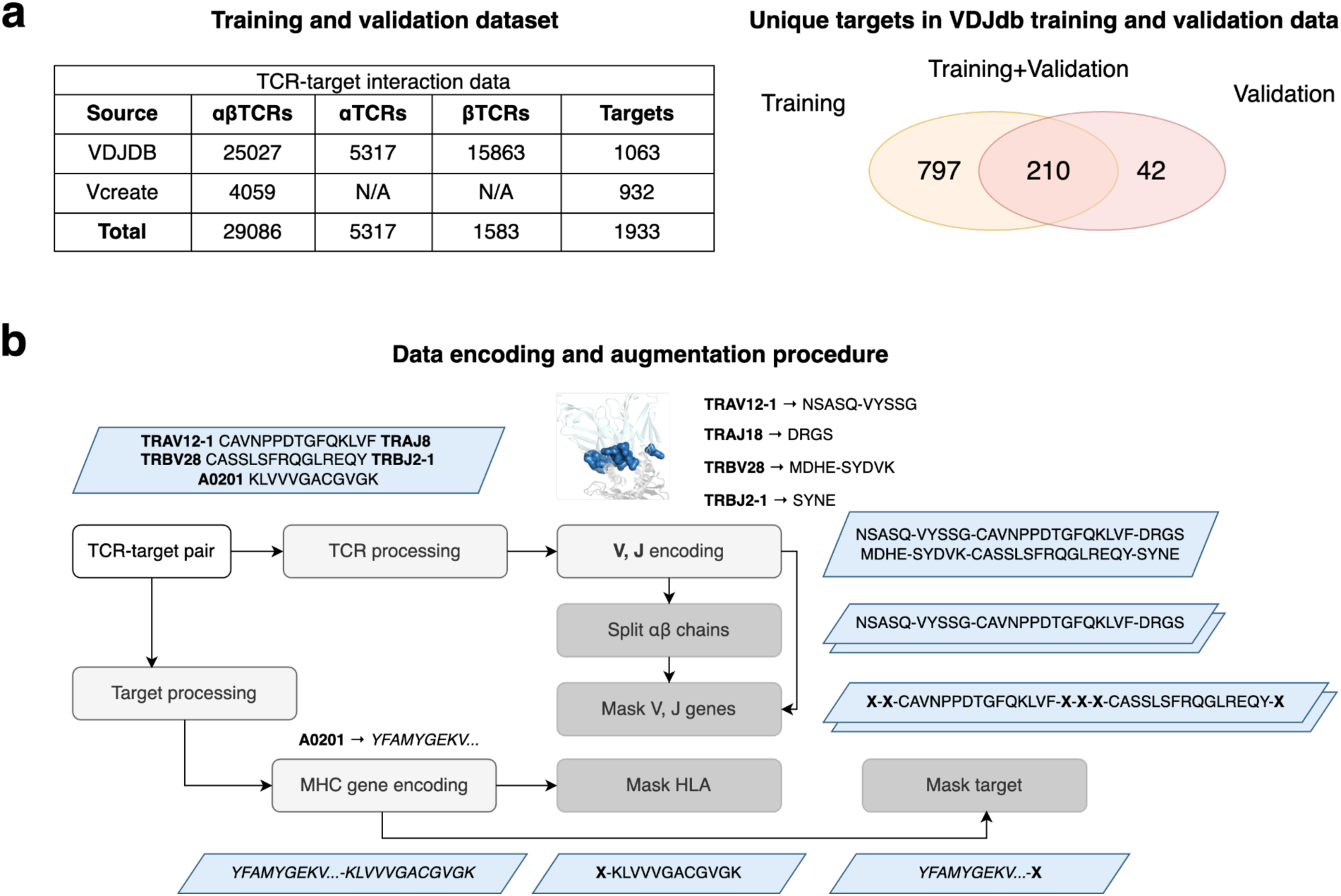
TAPIR training data and data augmentation. (**a**) TAPIR models are trained on larger and more diverse data than prior models, including more than 50k receptor sequences from VDJdb and proprietary Vcreate data paired with nearly 2000 targets. Our validation set includes 42 targets that models never encounter during training. (**b**) TAPIR learns from an augmented dataset that includes both paired and unpaired T-cell receptors, masked V and J genes, and masked alleles and target sequences during training. The data augmentation pipeline first processes V, J, CDR3, and MHC genes to represent TCRs and targets as sequences of amino acids. Additional training examples are then created following the steps above, for example “splitting” paired TCRs to create two new single chain training examples, or creating two new examples with masked V and J gene sequences. This data augmentation procedure allows TAPIR to learn how to reason about sequence inputs with different combinations of components (e.g., paired chain with CDR3 only, unpaired beta chain only with V, J, CDR3, etc.) and improves validation performance on novel targets.

**Figure 3:**
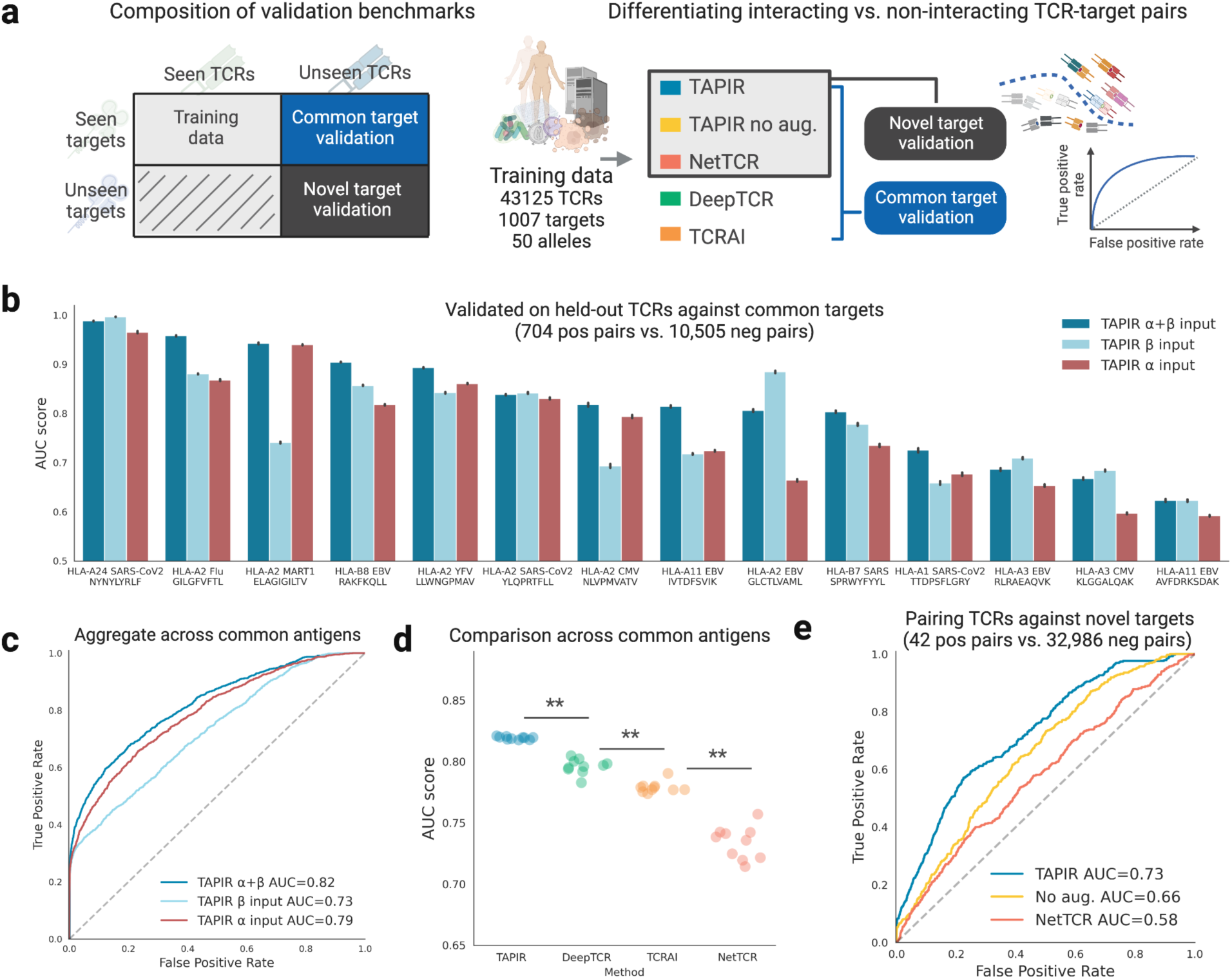
TAPIR prediction performance on common and novel antigen targets. (**a**) We compared TAPIR with several other models on benchmarks for common and novel targets, evaluating their ability to discriminate interacting vs. non-interacting TCRs via AUC. Models were trained on the same snapshot of VDJdb and performance was validated using held out data, including 15 common and 42 novel targets. (**b**) TAPIR model performance when given full TCR sequence input, just just a-chain, or just b-chain. (**c**) AUROC curves for TAPIR given full input, just alpha chain, or just beta chain. Providing complete TCR information (both alpha and beta chain) outperformed providing only alpha chain or beta chain (p<1e-5). (**d**) Comparison of 3 other leading models with TAPIR on common targets (**, p<1e-5). Two of these methods, DeepTCR and TCRAI do not encode target sequence information and so cannot be used to predict interactivity against novel targets. (**e**) AUC curves for the TAPIR model, an independent model trained on the TAPIR architecture where the data augmentation step was removed (“No aug.”), and NetTCR against novel targets.

**Figure 4:**
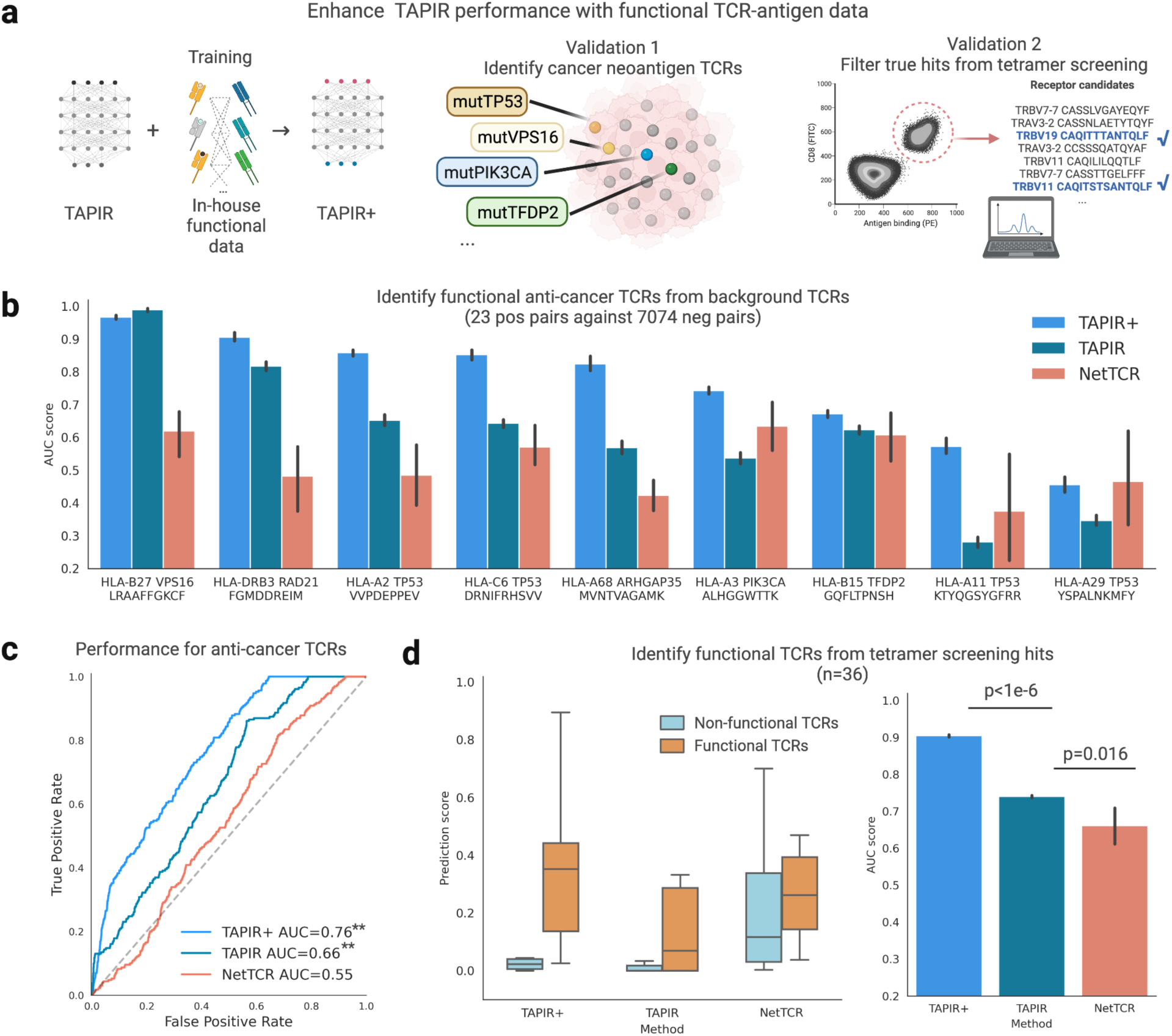
TAPIR trained with additional functional TCR-target interaction data performs better when predicting novel cancer targets and activating hits in tetramer screens. (**a**) We trained a new model (TAPIR+) on both public data and a proprietary dataset of 4059 TCRs with functional data against 932 targets. We then benchmarked TAPIR+ against two other models (TAPIR and NetTCR) trained only on identifying cancer neoantigen TCRs and filtering true hits from tetramer screens (**b**) Model performance in AUC for 9 novel cancer neoantigens that do not appear in model training data and which have been validated for functional activation with 24 TCRs. (**c**) AUC curves for models when discriminating TCRs associated with these targets among a background distribution of TCRs associated with 200+ alternative targets (**, p-val<1e-5). (**d**) Models were tasked with discriminating 13 activating TCRs from 23 binding but non-activating TCRs in tetramer screening data. On the left, we chart the range of scores for functional and non-functional TCRs. On the right, we plot model AUC.

In a series of analyses, we demonstrate the robustness and versatility of TAPIR. First, we retrain three previous state-of-art TCR prediction algorithms^18,19,23^ on a new, larger dataset, comparing TAPIR to these retrained methods on a benchmark of common antigen targets. We then demonstrate TAPIR’s capability to predict TCR interactions against novel targets on a dataset of held-out interaction data and examine how our in-house functional TCR-antigen dataset improves model performance for identifying functional antigen-specific TCRs (**Fig. 4**). We apply TAPIR to TCR repertoires from cancer patients carrying PIK3CA mutations and discover a novel and potent TCR against a common PIK3CA cancer driver mutation (**Fig. 5**). Finally, we show how the expressiveness of TAPIR can power other immune-related tasks, including predicting interacting MHC alleles from TCR sequences directly (**Fig. 6**), identifying key peptides in a TCR, and computationally generating antigen specific TCRs (**Fig. 7**).

**Figure 5:**
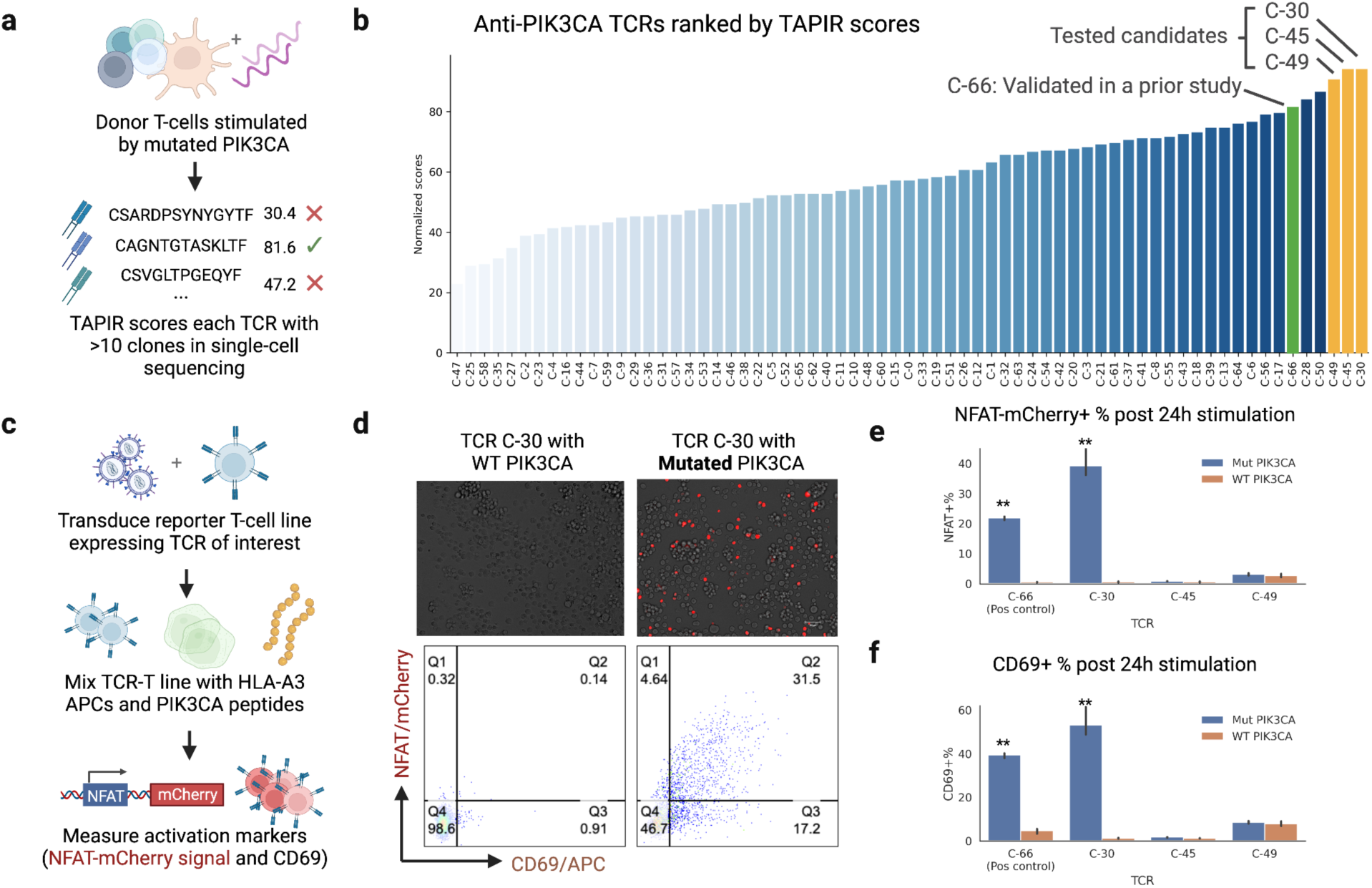
Identifying a novel TCR against mutated PIK3CA. When screening for TCRs against a mutated PIK3CA antigen target (A**L**HGGWTTK), TAPIR model scores help identify a novel TCR that we validated for function using NFAT and CD69 activation assays. (**a**) Donor T-cells were stimulated and expanded with mutated PIK3CA presenting autologous antigen presenting cells. T-cells were then sequenced with 10x Genomics single-cell sequencing. (**b**) TCRs with >10 clones in the screening were scored and ranked by TAPIR, and the three highest ranked TCRs were tested, along with a positive control TCR (C-66) from the previous study. (**c**) A reported T-cell line was transduced with the TCRs of interest and then mixed with HLA-A3 positive K562 cells and either wild type or mutated PIK3CA peptides. The reporter cell line expresses mCherry protein when the nuclear factor of activated T-cells (NFAT) is turned on. Activation markers NFAT and CD69 were measured 24h after cell mixing. (**d**) The highest TAPIR ranked candidate, C-30, was positive for NFAT and CD69 against mutated PIK3CA and not against the WT control. (**e**) The novel TCR C-30 demonstrated a stronger antigen-specific activation signal and less off-target effects than the positive control TCR C-66 (**, p<0.001). The other two candidates we tested, C-45 and C-49, do not show target-specific activation.

**Figure 6:**
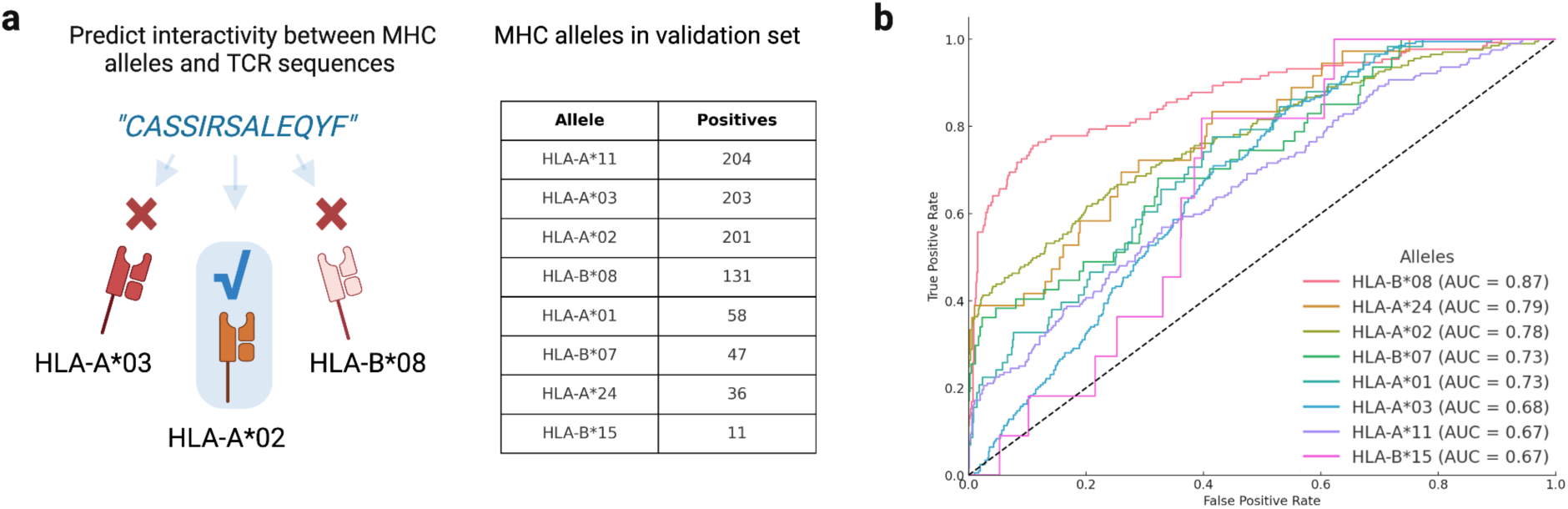
Predicting target MHC alleles from TCR sequences. (**a**) TAPIR’s data augmentation procedure allows the model to directly predict TCR interactivity against MHC alleles without requiring specific target information. We evaluate TAPIR’s performance on this task for 8 MHC alleles with at least 10 positive TCR examples in the validation set. (**b**) AUC curves and aggregate AUC scores performance are presented for these 8 alleles.

**Fig. 7.**
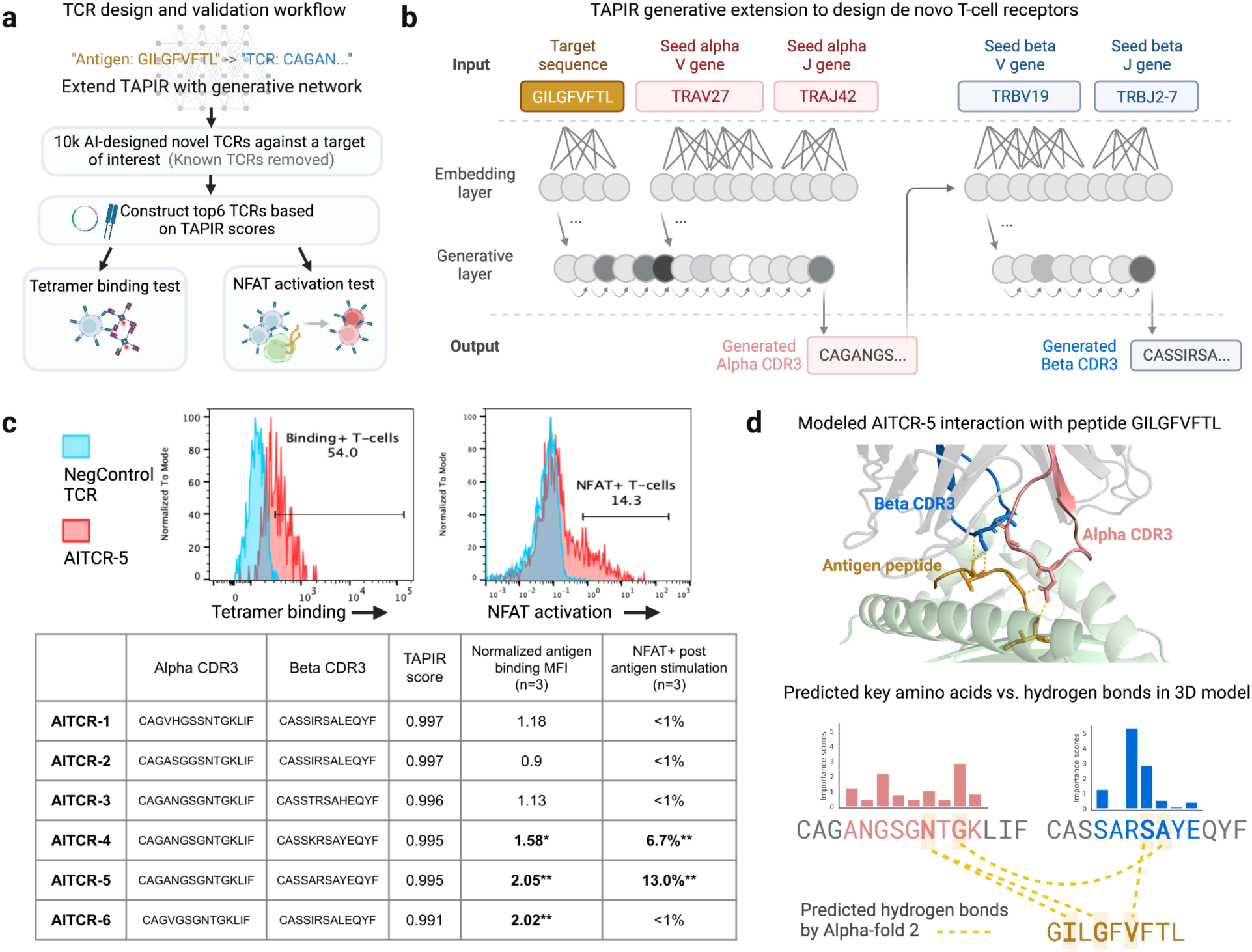
Extending TAPIR to design functional TCRs. (**a**) We extended TAPIR with a generative component capable of producing TCR sequences specific to target antigens. We then queried the extended model to sample and score 10,000 paired TCR alpha and beta CDR3 sequences against HLA-A2 presented influenza antigen peptide GILGFVFTL. Six TCR designs with the highest TAPIR scores were constructed and transduced into the TCR negative Jurkat NFAT-reporter line. Each TCR was evaluated on its ability to bind to HLA-A2-GILGFVFTL tetramer complex and to elicit T-cell activation signals (NFAT) when stimulated by GILGFVFTL-presenting antigen presenting cells (T2 cells). (**b**) The generative extension to TAPIR learns to predict CDR3 regions for alpha and beta TCRs given a target of interest and a starting set of V and J genes. First the alpha CDR3 region is predicted given the target and V and J genes, then the beta CDR3 region is predicted given the alpha chain, target sequence, and V and J gene set. Following TCR sequence generation, the full sequence is scored with the downstream original TAPIR model (**c**) TAPIR scores, normalized binding MFI, and T-cell activation results of six AI-designed TCRs against A2-presented GILGFVFTL. Tetramer binding MFI (median fluorescence intensity) was normalized to negative control TCR tetramer binding MFI. T-cell activation NFAT positive gate was set based on the top 1% of NFAT values in the negative control TCR group. Three AITCRs showed statistically significant binding to the target antigen (n=3, * p<0.05, ** p<0.001). Two AITCRs activated Jurkat cells when interacting with GILGFVFTL-presenting cells (n=3, ** p<0.001). (**d**) Correlation between predicted key amino acids with modeled TCR-antigen interactions for AITCR-5. The co-crystal structure of AITCR-5 and A2-presented GILGFVFTL antigen was modeled by AlphaFold 2 (red = alpha CDR3, blue = blue CDR3, orange = antigen peptide, green = HLA-A2, yellow dash line = predicted hydrogen bond). Amino acid importance scores for the binding regions of the CDR3s were computed with TAPIR. We highlight key amino acids (high scores) involved in the TCR-antigen and TCR alpha-beta hydrogen bond interactions (yellow dashed lines).

## Results

### Diverse TCR-antigen interaction data against hundreds of novel targets

The absence of sufficiently large and diverse training data is a key factor that has held back computational models for TCR-antigen prediction, making it difficult for models to generalize to new targets. In this paper, we leverage new datasets and data augmentation methods to increase the number of unique TCR sequences and antigen targets from which a model can learn.

We begin with a dataset of labeled human TCR sequence data hosted by VDJdb^22^. This dataset is generated by more than 200 studies and contains 46,207 unique human TCRs paired with 1063 pMHC targets across 57 MHC alleles. To our knowledge, prior machine learning models developed for TCR-antigen prediction have used either paired alpha-beta TCR sequences, or else only beta-chain sequences, and few models previously trained on targets with fewer than 10 known positive examples^7,20,24–26^. In contrast, models trained using our framework learn from both paired and single chain data, as well as a long tail of targets with few known positives, expanding the number of unique TCR sequences and targets observed in training (**Fig. 2a**). One key contribution of this paper is also a data augmentation procedure that expands TCR-target pair examples by 6 folds with several transformations (“masking”, **Fig. 2b**). Our procedure expands paired TCR sequences to create new single chain example sequences, generates additional training sequences with masked V and J genes (masking replaces a component of the sequence with a dedicated mask token “X”), and similarly expands pMHC target data with additional examples that mask the target sequence or MHC allele.

In addition to VDJdb binding data, we use a proprietary dataset of TCR-antigen functional associations. This data was generated as part of Vcreate in-house TCR discovery platform. Briefly, antigen presenting cells expressing a single target (pMHC single chain trimer^27^) are tagged with target DNA barcodes on their cell membrane, and T-cells with interacting TCRs are activated by these engineered antigen presenting cells and obtain target DNA barcodes via macrocytosis^28^ or trogocytosis^29,30^. TCR-antigen pairs are read out by performing single-cell sequencing on activated T-cells. This functional training set consists of 4059 paired alpha-beta TCR clones connected with 932 antigen targets (**Fig. 2a**). Antigen targets cover a variety of domains including cancer, autoimmune and infectious disease targets. Of these targets, 870 have no existing TCRs reported in public databases. Unlike the majority of data in public databases, which is based on pMHC multimer binding, all the TCR-antigen pairs in this dataset are associated through TCR activation. When training on this data, we again apply the data augmentation procedure (**Fig. 2b**).

### A generalized immune language model for TCR-antigen prediction

We developed the TAPIR architecture with two primary goals: first, to make predictions against targets never observed in training, such as cancer-associated neoantigens; and second, to maximize the signal that can be extracted from existing paired TCR-antigen data, which is noisy and collected in both single-chain and paired chain formats.

To make predictions against any target antigen, we draw inspiration from language models, which have demonstrated generalizability across diverse tasks and can be flexibly “programmed” with their inputs^31–34^. We experimented with many architectures when designing TAPIR and found that a two-tower architecture composed of independent CNN-based encoders for TCR and target sequences showed the best performance. By setting the target encoder’s input to a specific amino acid sequence (e.g., “KLVVGAVGV” produced by mutated cancer gene KRAS), a researcher can “program” TAPIR to produce TCR interactivity predictions for any target of interest, even those very different from what the model encountered during training (**Fig. 1**). TAPIR accepts target inputs as a combination of their amino acid sequences and the MHC allele by which they are presented, either of which can be missing. TCR sequences are similarly encoded using their CDR3 sequences and V and J genes, in paired or unpaired format. After the network encodes representations of a TCR and a target, these representations are combined and passed through a fully connected layer, then the network outputs a score between 0 and 1 indicating the likelihood of input TCR and target interacting (**Fig. 1**).

To better exploit the value of existing training data, TAPIR supports a flexible representation format for both TCRs and target sequences and learns from several tasks concurrently during training following the previously described data augmentation procedure (**Fig 2b**). After data augmentation, negative examples are created by repeatedly shuffling the combined pool of encoded TCR sequences to match with new targets. This is a common approach when predicting interactivity between sequences^18–20,23^ and ensures the distribution of TCR and targets is the same between positive and negative examples, which is important to avoid TCR or target specific bias.

We compared TAPIR to three previously published models from the literature: TCRAI, DeepTCR, and NetTCR^18,19,23^. Prior models are only able to handle a small number of targets (4-16), so we retrained all models on the same snapshot of VDJdb dataset (May 2023). From this snapshot we randomly sampled 90% of TCRs paired with antigen targets for training. The remaining 10% of the dataset of paired TCRs was used for validation, where we selected the 14 most common targets as a benchmark set and randomly selected up to 100 positive examples for each target (**Fig. 3a**). TCR examples in the validation set are strongly differentiated from TCRs in the training set when analyzed for similarity using the amino acid edit distance metric (median shortest edit distance between training and validation = 15, **Supplementary Fig. 1**). For TAPIR, we benchmarked performance when providing inputs as paired TCR sequences vs. single-chain TCR sequences (**Fig. 3b, 3c**). We noticed paired TCR inputs (no missing info) provided moderately better prediction accuracy but observed a few cases in which unpaired alpha or beta chain sequences offer comparable predictive performance. For example, providing the TCR alpha chain only performed similarly for predicting a famous melanoma antigen MART1 (ELAGIGILTV, **Fig. 3b**).

Comparing TAPIR with other methods trained on the same training set, the area under the curve (AUC) metric indicates that TAPIR, DeepTCR, and TCRAI all offer strong performance on common targets, with TAPIR reporting higher and more consistent scores (p-value<1e-5, **Fig 3d**). Notably, NetTCR, like TAPIR, is a general model that can take any antigen target as an input, whereas DeepTCR and TCRAI adopt categorical classes for each antigen and cannot be applied to targets outside of the training set. We next applied the two models capable of predicting TCR interactivity against novel targets – TAPIR and NetTCR – to a harder zero-shot benchmark task. We could not include TCRAI and DeepTCR in this benchmark as they cannot make predictions against novel targets. We selected all validation examples in the validation set that do not appear within >=2 edit distance from any target in the training set, leaving 42 novel targets (**Fig. 2a**). We also measured performance of the model against these targets as well as a smaller group of 10 targets with >0 evidence score in VDJdb. We observed much stronger performance for TAPIR with data augmentation both on the full set of novel targets (0.73 vs 0.65 AUC, **Fig. 3e**) and also on the smaller set of targets with non-zero evidence scores (0.78 vs 0.68 AUC). TAPIR significantly outperforms NetTCR (p<1e-5) despite training on the same data.

We also analyzed TCRs in the zero-shot validation benchmark task to compare them with the closest corresponding TCRs in the training set (**Supplementary Table 3b**). Nearly all of the TCRs (37 of 42) in this validation set are quite different from any TCRs in the training set, with the closest exhibiting a minimum edit distance of 12 edits. The remaining 5 TCRs have 0 edit distance to TCRs in our training set but are paired with different targets, and literature review confirms they are indeed cross-reactive TCRs ^8,35–38^. For example, two validation pairs are involved with a TCR with a beta CDR3 of CASSLWEKLAKNIQYF, which was determined to react with a bacteria protein fragment (MVWGPDPLYV, training set)^36^. The validation set included two targets that tested the model’s ability to match that TCR against human proinsulin variants (self-antigens RQWGPDPAAV and RQFGPDFPTI). The model achieves >0.90 AUC associating the correct TCR with these two self-antigen targets.

### Functional data helps models better predict TCR activation

While most existing TCR-target data captures binding relationships, TCR-target interaction is better defined by functional events associated with TCR activation, such as signal transduction, cytokine release, and killing. We next examined the additive impact of training TAPIR on a proprietary functional dataset that maps associations between 4059 TCRs and 932 targets, where 870 of these targets have no associated TCRs in VDJDB. We trained a new TAPIR model, TAPIR+, on both the VDJ snapshot and this new functional data and compared it to a baseline TAPIR and NetTCR model trained only on the VDJ snapshot.

To evaluate the impact of this functional data we constructed two new benchmarks (**Fig 4a**) based on TCRs validated for activation against targets (not just binding, as is the case for most public data). The first benchmark consists of 23 TCRs validated for function against 9 cancer neoantigen targets with gold standard activation markers such as CD69 and interferon γ secretion, which we gathered from 6 previous studies^16,39–43^. All 10 targets are “novel” to the model and never observed in the training data (**Supplementary Table 4** and **5**). We observe that TAPIR+ trained on additional functional data give strong improvements on this benchmark (**Fig 4b, 4c**). Additional training on functional data improves overall AUC from 0.64 to 0.73 for TAPIR (p<0.001), with NetTCR giving 0.59 (**Fig. 4c**). Performance is also more consistent for TAPIR models in comparison to NetTCR, with less variance across targets (**Fig 4b**). TCRs in this validation set are strongly differentiated from the training dataset with TCR edit distances of more than 11 from any training TCRs (**Supplementary Table 4b**).

The second benchmark comes from a TCR screening study conducted by Spindler et al^44^, where they ran binding-based screening to identify TCRs against 5 targets and functionally tested their top binders (**Supplementary Table 6** and **7**, n=36). The original authors identified 36 potential binding TCRs after several rounds of binding enrichment, but only 13 of these TCRs (36%) were validated in their T-cell activation assay (CD69). For this benchmark, we test whether models can distinguish the activating TCRs from the binding screening hits. Of the 5 targets in this benchmark, two are common antigens with abundant model training data (NLVPMVATV and GLCTLVAML) and three are rare with limited training data (CLGGLLTMV, VLEETSVML, and KTWGQYWQV). Overall, models with additional training on functional data demonstrated a significant increase in aggregate AUC performance from 0.75 to 0.90 (TAPIR vs. TAPIR+, p<1e-6), with NetTCR giving 0.66 (**Fig. 4d**). At 50% recall, both the TAPIR and TAPIR+ models show precision of 87.5% while NetTCR shows precision of 41% (**Fig 4d**).

### TAPIR discovers a novel TCR against mutated PIK3CA

FDA approved the first TCR therapy against solid tumors in 2022^45^, and there is increasing interest in discovering more anti-cancer TCRs^9,41,43,46^. One important use case for TCR-antigen prediction models is computational drug discovery – guiding the identification of T-cell receptors against clinical valuable targets such as shared cancer neoantigens^17,47^. One such target is PIK3CA, a gene frequently mutated in breast, colon, and lung cancers. PIK3CA encodes the catalytic subunit of the phosphatidylinositol 3-kinase enzyme, essential for cancer survival. More than one third of breast cancer patients carry at least one PIK3CA mutation, and PIK3CA H1047L mutation is the fourth most common PIK3CA mutation^48^. PIK3CA’s peptide fragment ALHGGWTTK is well presented by HLA-A*03:01 allele, and TCRs against A*03:01 presented ALHGGWTTK showed strong anti-cancer properties in xenograft mouse models^40^.

We investigated whether TAPIR could identify novel T-cell receptors that activate against mutated PIK3CA (**Fig. 5a**) using a population of TCRs sequenced from three PIK3CA mutated cancer patients^40^. TCRs in this population were previously enriched against ALHGGWTTK target and sequenced with 10x Genomics platform. We used TAPIR to score the 67 TCRs with more than 10 clones in this population against ALHGGWTTK (**Fig. 5b**). We ranked TCRs using two scores: *raw score*, the direct output score of the TAPIR model, and *antigen percentile*, a measure of how an individual TCR’s score against ALHGGWTTK compares to scores for 200 other common, rare, and novel antigens. We noticed that one TCR (C-66) identified and validated in the previous study ranked 5th in this data using the antigen percentile ranking and used that as the final ranking (**Fig. 5b**).

We then selected the top 3 TCRs (C-30, 45, 49) from the ranking to grow and test for function against the mutated PIK3CA peptide ALHGGWTTK and the wild type control AHHGGWTTK. Each of these TCRs is novel with distinct V, J, and CDR3 regions from previously reported TCRs for ALHGGWTTK antigens^40^. We previously established a Jurkat-based reporter T-cell line where mCherry fluorescence protein expression is controlled by a nuclear factor of activated T-cells (NFAT) promoter (**Fig. 5c**). All TCRs were transduced into our reporter lines for TCR function testing. We mixed transduced T-cell lines with HLA-A*03:01 expressing K562 cells pulsed with either wild type or mutated PIK3CA peptides and measured the NFAT-mCherry and CD69 levels (**Fig. 5c, 5d**, **Supplementary Fig. 2**). The C-30 TCR showed significant NFAT-cherry and CD69 up-regulation with mutated PIK3CA peptides but not with wildtype peptides (>30 fold difference, p<0.001, **Fig. 5d**, **Supplementary Fig. 2**). Furthermore, the novel C-30 TCR had higher activation level than the positive control TCR C-66 in both NFAT and CD69 level post stimulation (p<0.05, **Fig. 5e, 5f**). This case study demonstrates how TAPIR can enable the selection and identification of therapeutic TCRs from highly noisy primary T-cell pools.

### TAPIR can predict interactivity between TCRs and MHC alleles

TAPIR is trained on examples that mask peptide sequences in pMHC complexes to help models better generalize across the training data. As a result, TAPIR is also capable of predicting TCR interactivity with MHC alleles of interest in the absence of specific peptide information. For example, this functionality could be used to filter for true positive TCR hits from allele-specific screening experiments (e.g. tetramer screening). To our knowledge, TAPIR is the first tool capable of directly predicting interacting MHC alleles from TCR sequences.

To quantify TAPIR’s performance when predicting TCR-allele interaction, we evaluated it on data for 8 alleles in VDJDB with at least 10 positive examples (**Fig 6a**). TAPIR reports an overall AUC of 0.81 on this task, with strongest performance for HLA-B*08 (0.875 AUC), HLA-A*24 (0.79 AUC) and HLA-A*02 (0.778 AUC). Even alleles with relatively few training examples provide some predictive performance (0.67-0.73 AUC, **Fig 6a**). The model has unexpectedly low performance for HLA-A*03 (0.68 AUC) given the abundance of of HLA-A*03 training data. However, more than 95% of HLA-A*03 training examples concentrated around the single target KLGGALQAK. This highlights the importance of data diversity for TCR-target prediction models.

### TCRs designed by TAPIR show functional activation

In-silico TCR design has the potential to change the paradigm of TCR discovery pipelines, allowing for the eventual identification of clinical relevant TCR candidates without slow wet lab experiments ^6^. To examine TAPIR’s ability to aid in this process, we used our model to design a novel TCR against a common influenza antigen, GILGFVFTL, and validated the computationally generated TCRs with both binding and T-cell activation assays.

TAPIR is a discriminative model which produces interaction scores, and so without modification does not output TCR sequences directly. However, we can transform TAPIR into a generative algorithm by combining it with an additional component that generates target-specific TCR sequences for the downstream model to score. We can then sample from the combined network and select candidates with the highest predicted interactivity scores (**Fig. 7a**). For TAPIR’s generative component, we trained a simple autoregressive, recurrent neural network that, starting from any set of V and J genes and a target antigen, learned to construct amino acids for the alpha and beta chain CDR3 regions (**Fig. 7b**). To reduce the size of the search space when sampling novel TCRs, we also used TAPIR to identify V and J gene combinations for alpha and beta chain TCRs most likely to interact with GILGFVFTL. This is possible because TAPIR can make predictions for TCR sequences with missing CDR3 regions. Based on TAPIR’s gene scores we choose TRAV27 and TRAJ37 for alpha chain and TRBV19 and TRBVJ2-7 for beta chain, then sampled 10,000 candidate TCRs, removing any TCRs with alpha or beta chains observed independently in prior experimental or training data. We chose the top 6 hits to synthesize based on TAPIR’s interactivity scores.

Similar to our PIK3CA TCR validation, the six computationally designed TCRs (AITCRs) were transduced into reporter T-cell lines via lentiviral constructs. We performed both tetramer staining and NFAT activation assays (Fig. 7a, 7c) to evaluate the specificity of each candidate. Three of the six AITCR candidates show specific binding to the target GILGFVFTL above irrelevant TCR control (**Fig. 7c,** p<0.001). Two TCRs (AITCR-4 and AITCR-5) elicit NFAT activation signal when stimulated with GILGFVFTL peptides (**Fig. 7c,** p<0.001 while all six AITCRs have little to none baseline tonic signal. This provides a proof of principle that the field can use computational tools to design TCR candidates for drug development as TCR-target prediction models continue to improve.

We further investigated what key amino acids drive TAPIR’s predictions when designing these TCRs. AITCR-5 is the most potent TCR designed based on the data shown above. We exhaustively mutated each amino acid in the binding region of AITCR-5 alpha and beta CDR3s (**Fig. 7d**) and derived amino acid importance scores. These scores highlight A4, G6, N9, and G11 for alpha CDR3, and R6 and S7 for beta CDR3.

Interestingly, the <u>RS</u>A motif on beta CDR3 has been previously reported for GILGFVFTL binding^49^. We modeled the co-crystal structure of TCR-pMHC for AITCR-5 with AlphaFold 2^50^ and identified key residues based on hydrogen bond interactions. Our structure analysis highlighted more than half of “important” amino acids identified by TAPIR are involved in hydrogen bond formations (N9, G11, S6, A7, **Fig 7d**).

## Discussion

In this paper, we present a deep learning algorithm, TAPIR, that can analyze and predict TCR interactions against any antigen target, even targets with few or no previously reported interacting partners. TAPIR models are trained using a data augmentation method that enables them to improve upon the state-of-the-art prediction performance for common antigen targets, make predictions given incomplete sequence information (e.g., single chain TCRs, missing V genes), and predict TCR interactivity against MHC alleles when specific target sequences are not available.

Today, TCR-pMHC screening assays such as tetramer staining are a critical tool used to identify clinically valuable TCRs that bind and activate against a target of interest, which can then be further developed into therapies^40,44,51,52^. Unfortunately, screening assays have high false positive rates, often exceeding 90%, leading to many slow and expensive validation experiments to discover an interacting TCR^53^. Computational tools such as TAPIR have the potential to reduce these false positive rates and become an essential component of the TCR discovery pipeline. For example, when analyzing a dataset of tetramer data published by Spindler et al.^44^, a TAPIR score cut-off of 0.37 eliminates >95% of false positive TCRs from the tetramer screens (**Fig. 4d**). Notably, most clinical screening pipelines are focused on rare antigen targets such as cancer neoantigens, where training data is not available. TAPIR is useful today in this context, as we illustrate in benchmarks and our case study for mutated PIK3CA. We have set up a public server that other researchers can use to analyze their screening data with TAPIR at https://vcreate.io/tapir. This service is free for non-commercial use and is compatible with a variety of TCR sequence inputs, including paired and unpaired TCRs, as well as the file formats produced by the Adaptive Biotechnology and 10X Genomics screening platforms.

Beyond screening assays, in-silico discovery processes have the potential to revolutionize the field of TCR based immunotherapies by enabling the algorithmic screening of TCR candidates^6,9,20,33,34^, potentially many orders of magnitude faster than even the most high-throughput assays. In this study, we present a case study of one such in-silico discovery process, using TAPIR to design several novel TCR candidates that interact with the common influenza antigen. It is important to note that this case study was focused on an antigen target with many previously identified TCRs; discovery would likely be much more challenging for a target with fewer known examples. Nonetheless, our work provides an existence proof that in-silico TCR discovery is possible for targets with sufficient existing data. We are interested in testing TAPIR’s capability for similar kinds of discovery against rare and novel targets.

The overwhelming majority of existing TCR-antigen interaction data is capturing binding relationships observed in tetramer or dextramer screens^21,22^. In this paper, we observe that training TAPIR with proprietary dataset that captures functional interactions with hundreds of new targets significantly improves the model’s capability to accurately predict interactivity in situations where TCR activation is important, such as which TCRs will validate functionally in a tetramer screen, or which TCRs activate against a set of cancer neoantigens. TAPIR is capable of strong performance when trained largely on binding data, but this suggests that generating larger datasets of functional TCR data may be important for many applications, such as the discovery of TCRs for cell therapies.

It is important to note that TAPIR has limitations. For example, while performance in aggregate for novel binding and functional targets is compelling, for some targets TAPIR scores may not provide useful guidance. This is to be expected for a difficult zero-shot prediction task with such limited training data. So while TAPIR’s performance is strong enough to aid in screening applications, for most targets it is likely not strong enough to replace conventional wet lab screening approaches and should be leveraged as a supplemental method. As we and others generate more data, we look forward to characterizing the performance of TAPIR on additional targets.

Our work provides a new set of benchmarks and state-of-the-art performance on the task of predicting TCR interaction with unseen targets. In doing so, we lay a foundation that allows for many possible improvements. One such improvement would be to more directly incorporate structural data into the model. While TCR-target structural data is even more limited than today’s binding data, protein folding models like AlphaFold^50^ offer the potential to increase the available structural data for training in ways that might prove powerful for discriminating TCR interactivity with novel targets. Another promising avenue is to further expand TCR-target interaction algorithms to train on MHC presentation data and antibody data as supplementary tasks. In the same way that our data augmentation procedure led to improvements in generalization to novel targets, it is possible that shared relationships across an even more diverse set of tasks may further improve a model’s generalizability. Finally, new methods for data generation are perhaps the most obvious way to improve TCR-target prediction algorithms. We have demonstrated the improvements made possible when training on one such dataset that nearly doubled the number of unique targets observed by the model in training and plan to further develop the assays underlying that data.

## Code and data availability

Training and validation data of publicly available TCR-antigen pairs are listed in the Supplementary Table 1-7. Sources of anti-cancer TCRs are listed in Supplementary Table 4. Lentivirus and Sleeping-beauty vector backbones can be accessible via VectorBuilder’s vector retrieval page: https://en.vectorbuilder.com/design/retrieve.html. The VDJdb dataset is available at https://vdjdb.cdr3.net/. TAPIR is available for non-commercial use at https://vcreate.io/tapir/, and documentation is available for users at https://vcreate.io/tapir/tutorial.

## Acknowledgement

This work was partially supported by NSF SBIR grant (NSF-2213253), NIH SBIR grant (NIGMS, R43-GM143955), and Astellas Future Innovator Prize. We thank Russ Altman, Lisa Wagar, Karan Raj Kathuria, Yinnian Feng, Xiang Zhao, and Kevin Fast for discussing the manuscript.

## Materials and Methods

### Curation of publicly available TCR-pMHC datasets

We downloaded a snapshot of VDJdb taken on May 23, 2023 and filtered on “Human” for the “Species” column^22^. We randomly selected 10% of the data to use as a validation dataset for benchmarking. The remaining 90% of the data was used as a training dataset. To construct the secondary validation set composed only of novel targets, we filtered the previously described validation set to remove any TCR-antigen pairs with antigen targets that appeared within Levenshtein distance <=2 of any antigen targets in the training dataset. We then further filtered the novel target validation dataset to remove any TCRs with “Confidence Score” = 0.

### Independent training and validation TCR datasets

For our tetramer screening analysis, we downloaded the tetramer screening and validation dataset as reported in the Table 1 fo the original study^44^. For our validation dataset of functional anti-cancer TCRs we downloaded a list of cancer associated TCRs as reported by TCR3d^21^. We then curated this list to only include TCRs associated with activation reported against specific peptide sequences in the original literature^16,39–43^ and removed any TCRs present in the training VDJdb snapshot. For the mutated PIK3CA analysis, we obtained single-cell sequencing data from the original study^40^.

### Proprietary dataset of functional TCR-pMHC interactions

We previously generated a dataset of 4059 TCRs functionally associated with 932 antigen targets through a proprietary screening assay. This assay is capable of screening T-cells against a library of HLA-A2 antigen presenting cells presenting 1000 different antigens. The antigen presenting cell library is engineered such that when a T-cell activates against an APC an antigen-specific barcode is transferred from the APC to the T-cell via either trogocytosis or phagocytosis (patent application PCT/US22/36841). Antigen barcodes and receptor sequences are then read with 10x Genomics single cell sequencing and paired to construct a dataset of TCR-pMHC interacting pairs. We filtered this dataset to include only cell barcodes that included alpha-chain and beta-chain TCRs, as well as antigen barcodes. We filtered data to remove records with TCR UMIs less than 2.

### MHC allele validation dataset and scoring

For each allele with more than two associated TCRs in the validation set, we produce AUC curves measuring the model’s ability to differentiate allele-associated vs non-associated TCRs. We use a common set of 900 TCRs chosen randomly from the validation dataset as negative examples for each allele after removing any TCRs associated with the allele under analysis. Allele scores are computed with the TAPIR model by providing the model the allele psuedo-sequence^54–58^ and marking the associated antigen as a missing component using the ‘X’ masking input feature.

### Plasmid construction and viral transduction

T-cell receptor sequences were constructed based on IMGT ^59^ reference sequences, and codons were optimized for human cell expression (Benchling codon optimizer). TCR alpha chain and beta chain were co-expressed on a single plasmid with a T2A cleavage linker^60^. TCR constructs were cloned either into a sleeping beauty plasmid (EF1a promoter) or a 3rd generation lentivirus plasmid (EF1a promoter) with VectorBuilder service. For lentivirus construct, mouse TCR constant regions to improve TCR expression level as described previously^61^. Similarly, WT CD8A and CD8B TCR were co-expressed on a VectorBuilder sleeping beauty plasmid with a T2A cleavage linker without codon optimization. Lentivirus were produced with HEK293T cells and Mirus TransIT Lentivirus System (Mirus, 6650) according to the manufacturer protocol. Lentivirus was filtered with 0.45um PES filter (NEST, 380211). Jurkat cells or primary cells were transduced with the lentivirus for stable CD8 or TCR expression with the presence of 6ug/mL of DEAE (Sigma, D9885) as previously described ^62^.

### T-cell line and antigen presenting cells

Wild type Jurkat cells (TIB-152), K562 cells (CCL-243), HEK293T cells (CRL-3216), and T2 (CRL-1992) cells were obtained from the ATCC. J76 cells, a TCR negative Jurkat cell clone, are a generous gift from Mark Davis Lab from Stanford. Jurkat, J76, K562 cells were maintained in RPMI 1640 media (Cytiva, SH30096.FS) supplemented with 10% FBS (Sigma, F0926) and 10uM Glutamax (Gibco, 35050061). T2 cells were maintained in DEME media (Cytiva, SH30243.FS) supplemented with 10% FBS and 10uM Glutamax. To established HLA-A*03:01 monoalleic antigen presenting cell line, WT K562 were electroporated with a sleeping beauty transposase plasmid (VectorBuilder pRP[Exp]-CMV>T7/SB100X) and a sleeping beauty transposon plasmid (VectorBuilder sleeping beauty backbone, VB230803-1359aaa) co-expressing HLA-A*03:01 gene, B2M gene, and mNeonGreen gene as described previously^63^. The plasmids were constructed with VectorBuilder service (vector ID: VB900088-2243bzq). The electroporation was performed with Lonza 4D unit (Protocol CL-120) and 2ug total plasmid DNA per 1e6 cells. HLA-A*03:01 positive cells were sorted with Sony cell sorter SH800S based on the surface HLA level and GFP level (WT K562 cells have none to low HLA levels).

### Reporter T-cell line production with sleeping beauty

To produce a stable Jurkat T-cell line for TCR activation testing, three plasmids were electroporated into either WT Jurkat cells or TCR negative J76 cells: sleeping beauty transposase plasmid (VectorBuilder pRP[Exp]-CMV>T7/SB100X), sleeping beauty transposon plasmid expressing WT CD8 (VectorBuilder sleeping beauty backbone, VB230803-1359aaa), sleeping beauty transposon plasmid expressing NFAT promoter mCherry cassette. The NFAT promoter mCherry was constructed as described previously, and contained 4 NFAT response elements, a minimal IL-2 promoter, and codon optimized mCherry red fluorescent protein gene. All plasmids were constructed using VectorBuilder. Electroporation was performed with Lonza 4D unit (Protocol CL-120, 2ug total DNA per 1e6 cells).

### Antigen MHC complex tetramer staining

APC-conjugated HLA-A2*02:01-GILGFVFTL recombinant tetramers were obtained from MBL (Woburn, MA, TB-0012-2). Tetramer staining was performed according to the manufacturer protocol. Briefly, 1e6 TCR modified Jurkat cells or WT Jurkat cells were washed and resuspended in 100ul of PBS with 0.1% HSA. Cells were blocked with 10ul of human FcX block (BioLegend, 422302) for 10 min at room temperature. Then 1ul of tetramer was added and cells were incubated at 4C for 30 minutes. Samples were washed 2 times prior to analysis using flow cytometer Biorad ZE5. Positive gate was set by 0.1% tetramer+ WT Jurkat cells. Flow cytometry analysis was performed in FlowJo 10.8.

### T-cell activation assay

To assess activations of a TCR against an antigen of interest, CD8+ TCR-expressing Jurkat/J76 reporter cells (described above) were mixed with either peptide pulsed T2 cells (HLA-A*02:01) or peptide pulsed HLA-A*03:01 positive K562 cells for 24h. Peptides were synthesized with Elim Bio (Hayward, CA), and the default peptide concentration was 1ug/mL. Otherwise specified, a NYESO peptide (SLLMWITQV) was used as a negative control peptide for all activation assays to determine positive gate. CD69 and mCherry levels of TCR positive cells were analyzed with flow cytometry (Miltenyi MACSQuant Analyzer 16, see the gating strategy in Supplementary Fig. XXX). CD69 levels were measured with APC-conjugated anti-human CD69 antibodies (Biolegend, 310909, clone FN50). TCR levels were measured with BV421-conjugated anti-human TCR antibodies (Biolegend, 306722, clone IP26) or anti-mouse TCR antibodies (Biolegend, 109230, clone H57-597).

### TCR-pMHC co-crystal structure modeling

TCR-pMHC co-crystal structures were modeled with Alphafold2^50^ and ColabFold environment^64^. Briefly, variable regions of TCRs, HLA extracellular domain, antigen peptide, and B2M protein sequences were input into the ColabFold environment and separated by colon symbols. The modeling was conducted using default setting except the “relaxation” option was turned on. T100 GPU was used to accelerate the computing speed, and the structure of the highest ipTM score was analyzed for TCR-pMHC interactions. Hydrogen bond interactions were predicted with Pymol using 3.3 A as the cut-off^65^.

### Statistical test

Unless otherwise stated, statistically significant differences between distributions were determined by two-tailed paired student t-tests. We determined the statistical significance difference between two AUC curves (for example, Fig. 2d) using the fast DeLong test^66^. Any statistical P values below 1e-5 were denoted as P <1e-5 or P< 1×10-5.

### TCR sequence representation for TAPIR

We represent TCR sequences via the amino acids that constitute their variable CDRs. In particular, each TCR chain is: V_pseudo + CDR3 + J_pseudo, where V_pseudo and J_psuedo are shot “pseudo-sequences” of amino acids that under prior computational analysis were determined as most important for interaction (e.g., TRAV12-1 is NSASQ-VYSSG), and the “+” sign separates components with a “-” character. Alpha chain and beta chain TCRs are then combined as: aTCR + bTCR, again separating with “-”. For instance, a TCR under this representation can be expressed as:

### NSASQ-VYSSG-CAVNPPDTGFQKLVF-DRGS-MDHE-SYDVK-CASSLSFRQGLREQY-SYNE

Sometimes TCRs may not have both alpha and beta chains available or may be represented with missing components (see the following data augmentation section). In these cases, missing components are replaced with an “X” character. For example, a TCR with only an alpha chain is NSASQ-VYSSG-CAVNPPDTGFQKLVF-X-X-X-X. Or a TCR with only CDR3: X-X-CAVNPPDTGFQKLVF-X-X-X-CASSLSFRQGLREQY-X.

### Antigen and MHC sequence representation for TAPIR

Similar to TCRs, we represent pMHC complexes via the amino acids that represent the key regions for interaction, in particular: PEPTIDE + ALLELE_pseudo, where PEPTIDE is the short protein fragment known to present by MHC, and ALLELE_psuedo is a pseudo-sequence of amino acids describing the key structural regions of the MHC allele, once more separated by “-”. As with TCRs, either component can be missing and replaced with “X”. Several examples:

- YFAMYGEKVAHTHVDTLYVRYHYYTWAVLAYTWY-KLVVVGACGVGK
- X-KLVVVGACGVGK (antigen only)
- YFAMYGEKVAHTHVDTLYVRYHYYTWAVLAYTWY-X (MHC allele only)

### Augmentation procedure for TCR-pMHC datasets

Data augmentation expands a set of positive training examples by generating additional positive examples with various components of TCRs and pMHC sequences masked. Specifically, for each (TCR, pMHC) pair in a dataset of positives, new examples are generated and added to the set of positives for training with the following components masked: beta-chain TCR, alpha-chain TCR, V and J genes, antigen target, and HLA-allele. As described in the previous section on representation, masked components are replaced with an ‘X’ token to indicate a missing component.

### TAPIR neural network architecture

A TAPIR model takes TCRs and pMHC complexes of interest as inputs of amino acid sequences and predicts their likelihood of interaction. The TCR and pMHC sequences are embedded using a vector representation learned by the model during training, then encoded via independent multi-layer convolutional encoders into vectors. These vectors are concatenated and passed through a fully connected dense layer, then a binary classification layer that gives a final predicted interaction score between 0 and 1.

#### Embedding amino acid sequences

the model takes input sequences of amino acids and embeds them into a vector used by downstream model components. First each character in the vocabulary of amino acids (with the addition of “-” and “X”) is converted into a number, which an embedding layer then maps onto a learned embedding. The dimensionality of the embedding used in our paper is 64.

#### Convolutional encoders

TCR and pMHC sequences are then processed through independent convolutional encoders. These encoders transform an amino acid sequence into a vector through three convolutions with batch normalization and pooling. Each layer performs a 1D convolution (with kernel size=3, stride=1, and padding=1) over the sequence to produce a new, deeper output channel dimension. Then values are batch normalized over the output channel dimension and processed with MaxPool. The final output matrix is flattened across the final output channels to produce a vector encoding for the sequence. For TCR sequences the output channel dimensions are D1=64, D2=128, D3=256 and for pMHC sequences they are D1=32, D2=64, D3=128.

#### Concatenation, dense layer, and classification

after TCR and pMHC sequences have been encoded, they are concatenated and processed through a fully-connected dense layer of 256 neurons. This 256-dimensional vector is then processed through binary classification module to produce an output score.

### TAPIR training process

Three TAPIR models are described in this paper: a model trained on VDJdb data with data augmentation, a model trained on VDJdb without data augmentation, and a model trained on both VDJdb and proprietary data with data augmentation. With the exception of the data augmentation step (which creates new positive examples, as described above), the training process is the same across all three model types.

First, negative training examples are created by repeatedly (3x) shuffling the pMHC targets associated with TCRs and labeling the resulting pairs as negative. Note that it is possible that randomly shuffling pMHC targets will sometimes result in correct pairings, but this is low probability and better than having different distributions of pMHC targets for the positive and negative examples, which could introduce bias into the model.

Models are trained for 20 epochs, shuffling the data between epochs, with a batch size of 256 and the Adam optimization function with learning rate=0.001, betas=(0.9, 0.999, epsilon=1e-08, and weight_decay=0. Loss is computed across each bach using a cross-entropy loss function and model weights are updated using backpropagation.

To further improve performance, we ensemble 64 TAPIR models for each model type, where scores are computed by the mean across the ensemble.

### Benchmarking TAPIR against other published methods

#### TCRAI

We used the scripts with default parameters as described by Zhang et al^23^ to train TCRAI models on our snapshot of VDJdb training data. TCRAI models are binary classifiers and do not take an antigen target as input, so this benchmark required training 14 models, one per antigen class. For each model, TCRs for one benchmark antigen were labeled as positives and TCRs associated with any other antigen were labeled as negative. TCRAI can not be trained with both single and paired TCR data, so we included only paired TCR data in training.

#### DeepTCR

We used provided scripts with default parameters for the latest version of DeepTCR^19^ to train models on our snapshot of VDJdb training data. As DeepTCR is a multi-class classifier, we labeled TCRs for each of the 14 common antigens in the VJDdb training snapshot with a class index. Models then learned to differentiate these classes in training. V and J genes are optional for DeepTCR, and we included them in training as this improved DeepTCR’s performance. DeepTCR does not support simultaneously learning from both paired and unpaired TCRs, so we only included paired TCR data in training.

#### NetTCR

Again we used provided scripts with default parameters to train NetTCR^18^ models on the VDJdb binding data snapshot. NetTCR, like TAPIR, is a general model that takes both TCR sequences and antigen sequences as input, so known binding pairs are labeled as positive. We generated negative training examples using the same shuffling method as TAPIR (which is also used in the NetTCR paper). NetTCR only supports peptides up to length 9, so peptides longer than this were shortened. NetTCR does not support V and J genes, so these were dropped from training and validation examples. Finally, NetTCR does not support training simultaneously on paired and unpaired TCR sequences, so we trained only on paired sequences.

For each model architecture, we trained and validated ten times with different random seeds to compute p-values, standard deviation, and variance in performance. Because TCRAI, DeepTCR, and NetTCR do not support simultaneous training on paired and unpaired TCR sequences, we evaluated all model architectures only on paired TCR sequences in the validation set for the benchmark.

### TCR generative model and top candidate selection

To transform TAPIR into a generative model, we trained a new generative neural network from which TCR sequences can be sampled and passed to the original TAPIR network for scoring. Concretely, we use a Long Short-Term Memory (LSTM) based network^67,68^ to generate CDR3 regions for alpha and beta chain T-cell receptors when given V and J genes for alpha and beta chain and a target antigen of interest. Training this network is broken into two tasks which occur sequentially: predicting the CDR3 regions for alpha chain, then predicting the CDR3 region for beta chain.

For predicting alpha chain CDR3 regions, the network is passed the concatenation of the target and alpha V and J genes. Encoding of the V and J genes and separation of components is performed as described previously. The network is then trained to predict each amino acid of the alpha CDR3 sequence until an end of sequence (EOS) token. For predicting beta chain CDR3 regions, the target and alpha chain sequence (including the previously generated alpha CDR3) is fed to the LSTM, along with beta V and J genes. The network then predicts beta CDR3 amino acids until an EOS token.

This component consists of a single layer LSTM with a hidden size of 256 connected with 0.4 dropout to a classifier over a space of 20 amino acids + the EOS token. We used a learning rate of 0.001 with Adam, a teacher forcing rate of 0.5 and trained on all public TCR-target data.

To sample from the generative version of TAPIR, the generative LSTM component is sampled with a temperature of 1 to produce a number of candidate TCR sequences (e.g. 10,000) for a given target and V/J gene set. The downstream TAPIR model then scores these sequences against the target for which they were generated (score 0-1). For the GILGFVFTL validation experiments, generated TCR sequences with the highest scores were further tested in T-cell assays.

### TCR amino acid importance score calculation

To estimate the importance of each amino acid in TCR CDR3 antigen regions, we created variants of CDR3 of interest with exhaustive single mutations in each position of interest. As the first three and last three amino acids are often not involved in antigen binding, these amino acids can be excluded from the analysis. TAPIR produced a prediction score for every variant. Absolute value of average delta of new scores minus the WT CDR3 were calculated.

