## Supplementary Figures for "TAPIR: a T-cell receptor language model for predicting rare and novel targets"

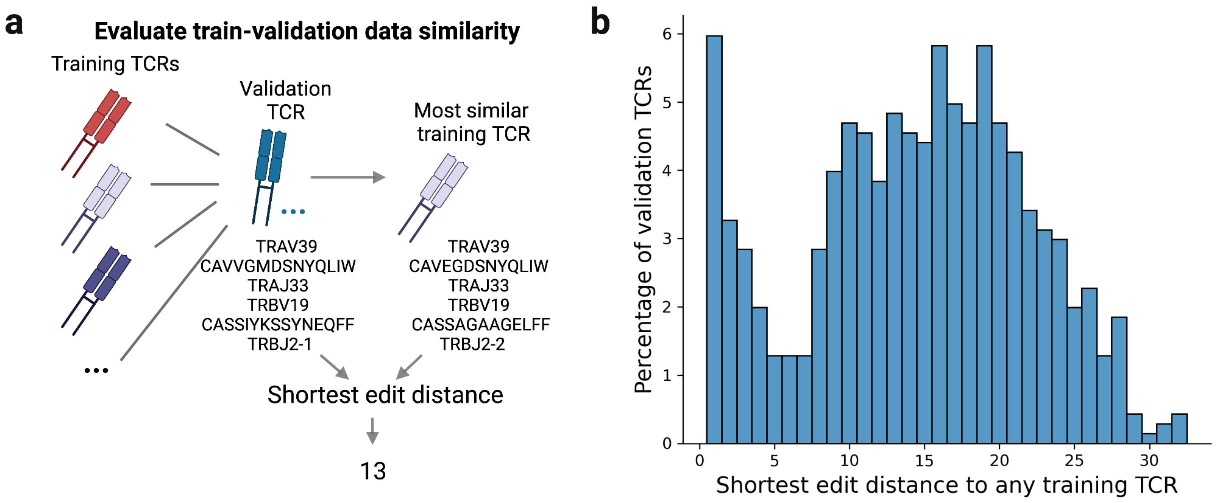


**Supplementary Figure 1. Train-validation data similarity analysis.** To further interrogate TAPIR's ability to generalize across TCRs and targets, we performed edit distance analyses on each of the held out validation sets. (**a**) Edit distance was computed using the Levenshtein edit distance method to find the closest TCR in the training set for each TCR in each of our validation sets. (**b**) We computed a histogram of these edit distances for the validation TCRs against 14 common antigen targets. For this dataset, 85% of the validation examples have an edit distance of more than 5 edits from the closest training example. No TCRs were shared (i.e., 0 edit distance) between training and validation. We also computed edit distances and reported the closest training TCRs and corresponding targets for all TCRs in our other two validation sets: a collection of novel targets from VDJdb (**Fig 3e**, **Supplementary Table 3b**) and a set of novel cancer-related targets with functional evidence (**Fig4b**, **Supplementary Table 4b**). With the exception of 5 cross-reactive TCRs in **Supplementary Table 3b**, all TCRs in these validation sets have more than 10 edits from the nearest TCRs in training data.

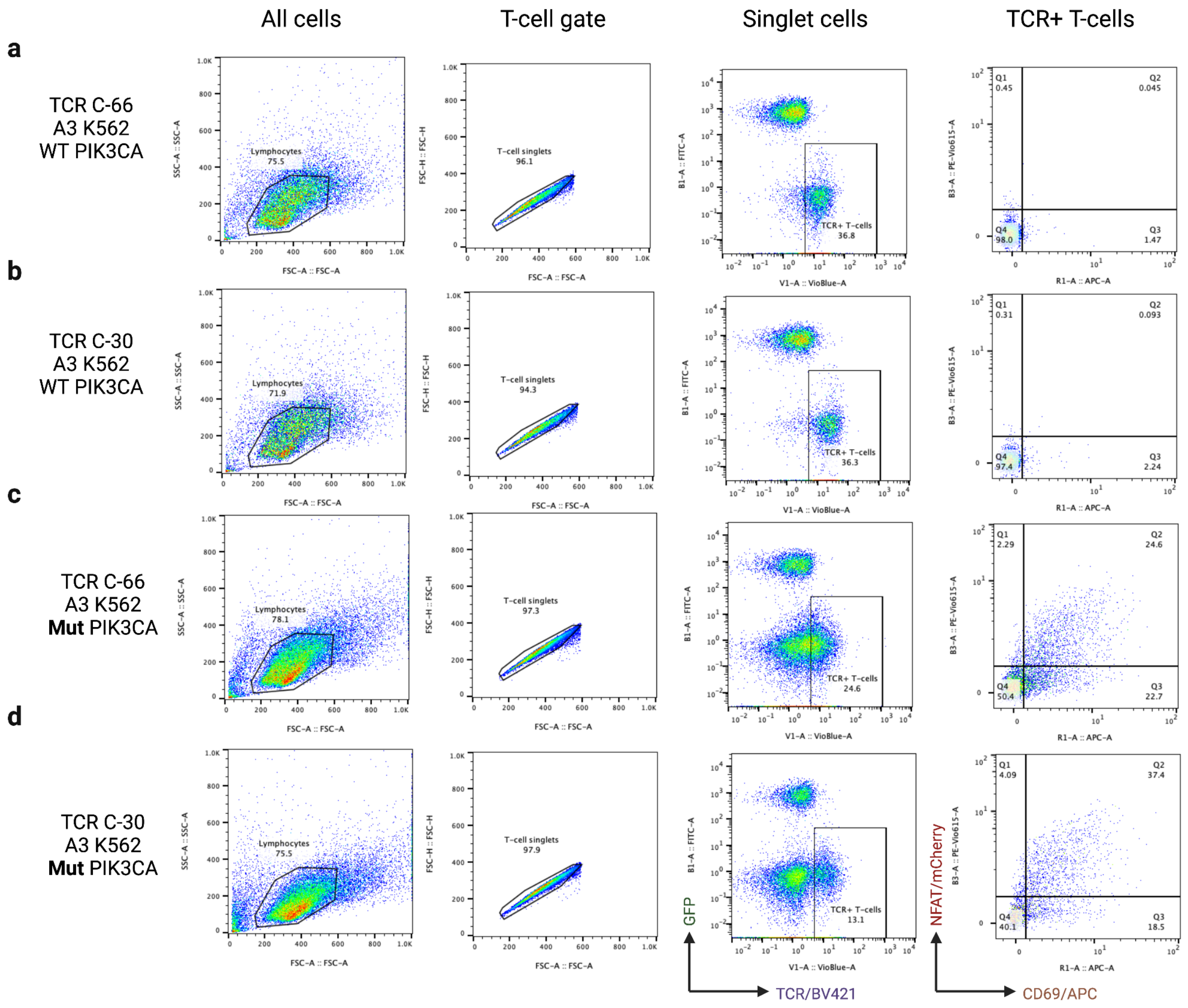


**Supplementary Figure 2. Flow cytometry gating strategy and representative data of anti-mutPIK3CA TCRs.** T-cells were first gated based on FSC-A and SSC-A for small lymphocyte size, and singlet cells were selected with the FSC-A and FSC-H gate. CD69 and NFAT-mCherry levels were measured on TCR+ and GFP low cell population (HLA-A3 K562 cells have bright GFP levels). Representative flow data were plotted for four co-incubation conditions: **a**) TCR C-66 T-cells with HLA-A3 K562 and WT PIK3CA peptide (AHHGGWTTK), **b**) TCR C-30 T-cells with HLA-A3 K562 and WT PIK3CA peptide, **c**) TCR C-66 T-cells with HLA-A3 K562 and mutated PIK3CA peptide, **d**) TCR C-30 T-cells with HLA-A3 K562 and mutated PIK3CA peptide (A**L**HGGWTTK). Cognate TCR-antigen interactions increased both NFAT-mCherry and CD69 levels for C-66 (positive control) and C-30 (identified in this study) TCR expressing T-cells. In addition, T-cell surface TCR levels decreased post cognate antigen stimulation (**c**,**d** compared to **a**,**b**), a hallmark of T-cell activation.
